# Transcriptomic evidence that von Economo neurons are regionally specialized extratelencephalic-projecting excitatory neurons

**DOI:** 10.1101/627505

**Authors:** Rebecca D Hodge, Jeremy A Miller, Mark Novotny, Brian E Kalmbach, Jonathan T Ting, Trygve E Bakken, Brian D Aevermann, Eliza R Barkan, Madeline L Berkowitz-Cerasano, Charles Cobbs, Francisco Diez-Fuertes, Song-Lin Ding, Jamison McCorrison, Nicholas J Schork, Soraya I Shehata, Kimberly A Smith, Susan M Sunkin, Danny N Tran, Pratap Venepally, Anna Marie Yanny, Frank J Steemers, John W Phillips, Amy Bernard, Christof Koch, Roger S Lasken, Richard H Scheuermann, Ed S Lein

## Abstract

von Economo neurons (VENs) are bipolar, spindle-shaped neurons restricted to layer 5 of human frontoinsula and anterior cingulate cortex that appear to be selectively vulnerable to neuropsychiatric and neurodegenerative diseases, although little is known about other VEN cellular phenotypes. Single nucleus RNA-sequencing of frontoinsula layer 5 identified a transcriptomically-defined cell cluster that contained VENs, but also fork cells and a subset of pyramidal neurons. Cross-species alignment of this cell cluster with a well-annotated mouse classification shows strong homology to extratelencephalic (ET) excitatory neurons that project to subcerebral targets. This cluster also shows strong homology to a putative ET cluster in human temporal cortex, but with a strikingly specific regional signature. Together these results predict VENs are a regionally distinctive type of ET neuron, and we additionally describe the first patch clamp recordings of VENs from neurosurgically-resected tissue that show distinctive intrinsic membrane properties relative to neighboring pyramidal neurons.

## Introduction

von Economo neurons (VENs) are a morphologically-defined neuron type with a large, characteristic spindle-shaped cell body, thick bipolar dendrites with limited branching and a moderate density of spines, and often an axon initial segment that emanates from the side of the cell body ^1,2,3^. VENs have been described in several large-brained mammals, such as humans, great apes, macaques, cetaceans, cows, and elephants, but not in rodents ^4,5,6,7,8,9,10,1,11^. In humans, they are restricted to the anterior cingulate (ACC), frontoinsular (FI), and medial frontopolar regions of cerebral cortex ^12^, while in most other species they are also found in the frontal and occipital poles ^13^ and may not be restricted to layer 5. Fork cells, another distinctive morphological-defined neuron type, are often found in the same brain regions as VENs and are similarly characterized by a single large basal dendrite, but differ from VENs by having a divided apical dendrite ^1,14^. VENs and fork cells appear to be selectively vulnerable neuron types, as loss of these cells has been observed in behavioral variant frontotemporal dementia (bvFTD) ^15,16,17^. Loss of VENs has also been observed in several neuropsychiatric disorders, including schizophrenia ^18^ and suicidal psychosis ^19^, as well as in autism ^20^, agenesis of the corpus callosum ^21^, and possibly Alzheimer’s disease ^22,23^.

Very little is known about VEN cellular phenotypes beyond their hallmark morphology, especially in human cortex. Human FI and ACC neurosurgical resections are extremely rare for functional studies, and VEN sparsity without some form of genetically-based labeling makes their analysis difficult. VENs have been described in rhesus monkey and tract tracing studies suggest that they might project to ipsilateral ACC and contralateral anterior insula ^4,24^, as well as to more distant targets in the parabrachial nucleus of pons and the midbrain periaqueductal gray ^25,26^. Molecular analyses of human VENs have been more fruitful since these techniques can be applied to postmortem human tissues. For example, a recent study using *in situ* hybridization (ISH) data from the Allen Brain Atlas identified *ADRA1A*, *GABRQ* and *VMAT2* as VEN marker genes ^27^, and a study using laser microdissection of VENs followed by RNA-sequencing identified additional potential VEN marker genes ^28^. VENs have also been reported to express serotonin receptor 2B (*HTR2B*) and dopamine receptor D3 (*DRD3*) ^29^, and the Schizophrenia-associated protein DISC1^4,25^. Additionally, they express transcription factors *FEZF2* and *CTIP2* ^26^ which are required for generating subcortical projection neurons in mice ^30^, and this has been used as evidence that VENs are subcortically-projecting neurons, referred to here as extratelencephalic-projecting excitatory neurons (ET) (though we acknowledge that ET neurons may not strictly project to subcortical structures and may have telencephalic collaterals ^31^). However, *Fezf2* is not specific for ET neurons but is also expressed in near-projecting pyramidal neurons in adult mouse ^32^, and expression of many cellular marker genes is not conserved between mouse and human ^33,34^. Furthermore, many of the reported markers of VENs are not exclusive to these cells but are also expressed in fork cells and pyramidal-shaped neurons. This highly incomplete characterization leaves unresolved many questions about whether morphologically-defined VENs represent a molecularly-distinct cell type and what their other properties are.

Single cell RNA-sequencing (scRNA-seq) has emerged as an effective strategy for classifying and characterizing cell types in complex brain tissues, and single nucleus (sn) RNA-seq can be used on frozen postmortem human brain specimens ^35,36^. Applied to cortex, this approach reveals a high degree of cellular diversity, with upwards of 100 transcriptomically-defined cell types in any cortical area ^33,32,37,38^. Furthermore, these data enable quantitative alignment of cell types across brain regions and between species to establish identity by transcriptional similarity using new computational strategies for mapping of transcriptomic types between datasets ^39,40,41^. Such alignment permits prediction of cellular properties and projection targets in human based on properties described in well-studied mouse cell types ^33^.

To reveal the transcriptomic signature and predict properties of VENs, we performed snRNA-seq on nuclei from layer 5 of FI and compared to similar data from mouse visual and human temporal cortex. We find a single transcriptomic cluster expressing several known markers for VENs that aligns with ET neurons in mouse cortex, as well as a putative transcriptomically-defined ET cluster in human temporal cortex that has a distinctive regional signature compared to FI. We identify many novel markers for this cluster and demonstrate that they are co-expressed in a combination of pyramidal neurons, VENs, and fork cells. Finally, we present a case study with the first electrophysiological recordings of putative VENs, and show that they have distinctive intrinsic membrane properties from neighboring layer 5 pyramidal neurons.

## Results

### Transcriptomic cell types in layer 5 of FI

We employed snRNA-seq ^35,36^ to profile nuclei from FI of two postmortem human brain specimens (Fig. 1a) as previously described ^42,33^. Briefly, layer 5 was microdissected from fluorescent Nissl-stained vibratome sections of FI and nuclei were liberated from tissue by Dounce homogenization. NeuN staining and fluorescence-activated cell sorting (FACS) were used to enrich for neuronal (NeuN+) and non-neuronal (NeuN-) nuclei (**Supplementary Figure 1a**). RNA-sequencing was carried out using Smart-seq2, Nextera XT, and HiSeq sequencing. In total 879 nuclei were processed for RNA-seq and were sequenced to a median of 4 million mapped reads. Median gene detection (expression > 0) was 10,339 for excitatory neurons, 9,426 for inhibitory neurons, and 6,146 for non-neuronal cells, consistent with previous reports ^42,33,42^ (**Supplementary Fig. 1b**).

**Figure 1.**
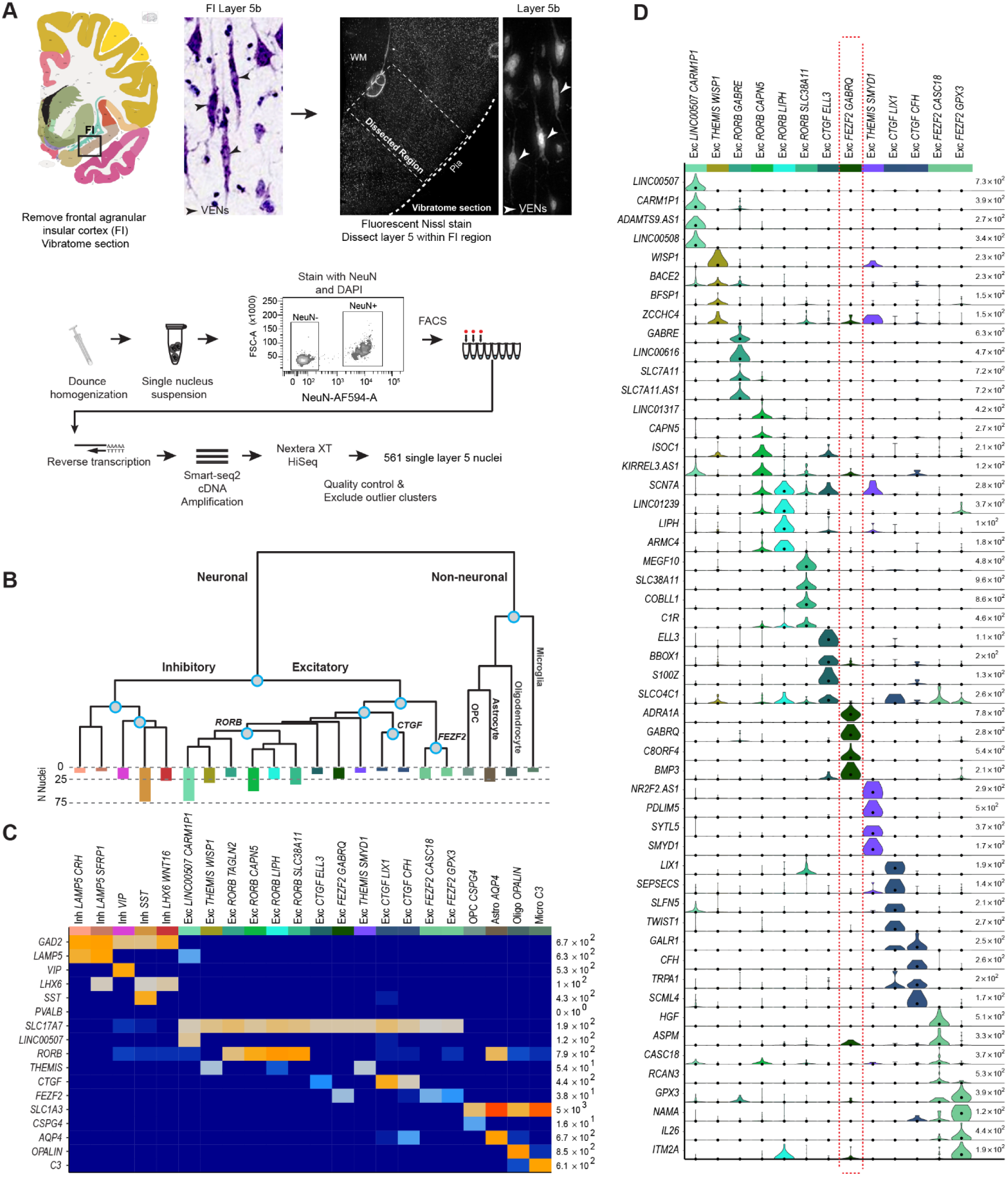
Cell type characterization in human frontal agranular insular cortex (FI). **(A)** Schematic diagram illustrating nuclei isolation from postmortem human brain specimens. The FI region was isolated, vibratome sectioned, stained with fluorescent Nissl, and layer 5 was dissected and processed for nuclei isolation, fluorescence-activated cell sorting (FACS), and RNA-sequencing. Examples of cells with morphologies typical of von Economo neurons (VENs) are shown in the images of Nissl-stained tissues (arrowheads). In total 561 single layer 5 neurons passed quality control. **(B)** Hierarchical representation of 18 neuronal (5 inhibitory, 13 excitatory) and 4 non-neuronal transcriptomic cell types based on median cluster expression. Major cell classes are labeled at branch points in the dendrogram. The bar plot below the dendrogram represents the number of nuclei within each cluster. Cluster-specific colors and labels are used in all subsequent figures. **(C)** Heatmap showing the expression of cell class marker genes across all clusters. Maximum expression values for each gene are listed on the far right-hand side of the plot. Gene expression values are quantified as counts per million of intronic plus exonic reads and displayed on a log10 scale. **(D)** Violin plots showing expression of four marker genes per excitatory cluster. Each row represents a gene, black dots show median gene expression within clusters, and maximum expression values for each gene are shown on the right-hand side of each row. Gene expression values are displayed on a linear scale.

Iterative clustering was used as described ^42,33,32^ to group nuclei by gene expression similarity. Briefly, high variance genes were identified while accounting for gene dropouts, expression dimensionality was reduced with principal components analysis (PCA), and nuclei were clustered using Jaccard-Louvain community detection. Clusters containing cells from only a single donor as well as nuclei mapping to low-quality outlier clusters (n=318) were excluded from further analysis, leaving a total of 561 high quality nuclei. We identified a robust set of 22 transcriptomically-defined clusters (Fig. 1b) that contained cells from both donors at roughly comparable proportions within broad classes (**Supplementary Fig. 1c, d**). Five inhibitory neuron types spanning all expected subclasses (two *LAMP5* types, *VIP*, SST, and a *LHX6*+/*SST*-cluster corresponding to *PVALB*), 13 excitatory neuron types, and four major non-neuronal cell types (oligodendrocyte precursor cells, oligodendrocytes, astrocytes, and microglia) were identified (Fig. 1c). Clusters were named using a broad class marker in combination with a highly specific marker, as described previously ^33^.

Excitatory clusters in FI expressed broad class markers previously identified in human middle temporal gyrus (MTG) (Fig. 1c, d) ^33^. One cluster had high expression of the MTG upper layer marker *LINC00507* and likely represents deep layer 3 pyramidal neurons sampled at the layer 3/5a boundary, since FI is agranular and does not contain layer 4. Three clusters express *CTGF*, a canonical marker for deep layer 6 neurons in mouse that has more widespread layer 6 expression in human ^34^, suggesting these clusters represent cells captured at the layer 5b/6 border. Two clusters highly express *THEMIS*, which is also expressed in layer 5 and 6 excitatory neuron types in MTG ^33^. Four clusters express *RORB*, which marks a subset of cells localized throughout layers 3-5 in MTG ^33^ and has a similar pattern of expression in FI (Fig. 2). Finally, we find 3 clusters with high expression of *FEZF2*, previously shown to be expressed in VENs ^26^, subcortically-projecting and near-projecting excitatory neurons in mouse cortex ^32^, and several deep layer excitatory types in human MTG ^33^.

**Figure 2.**
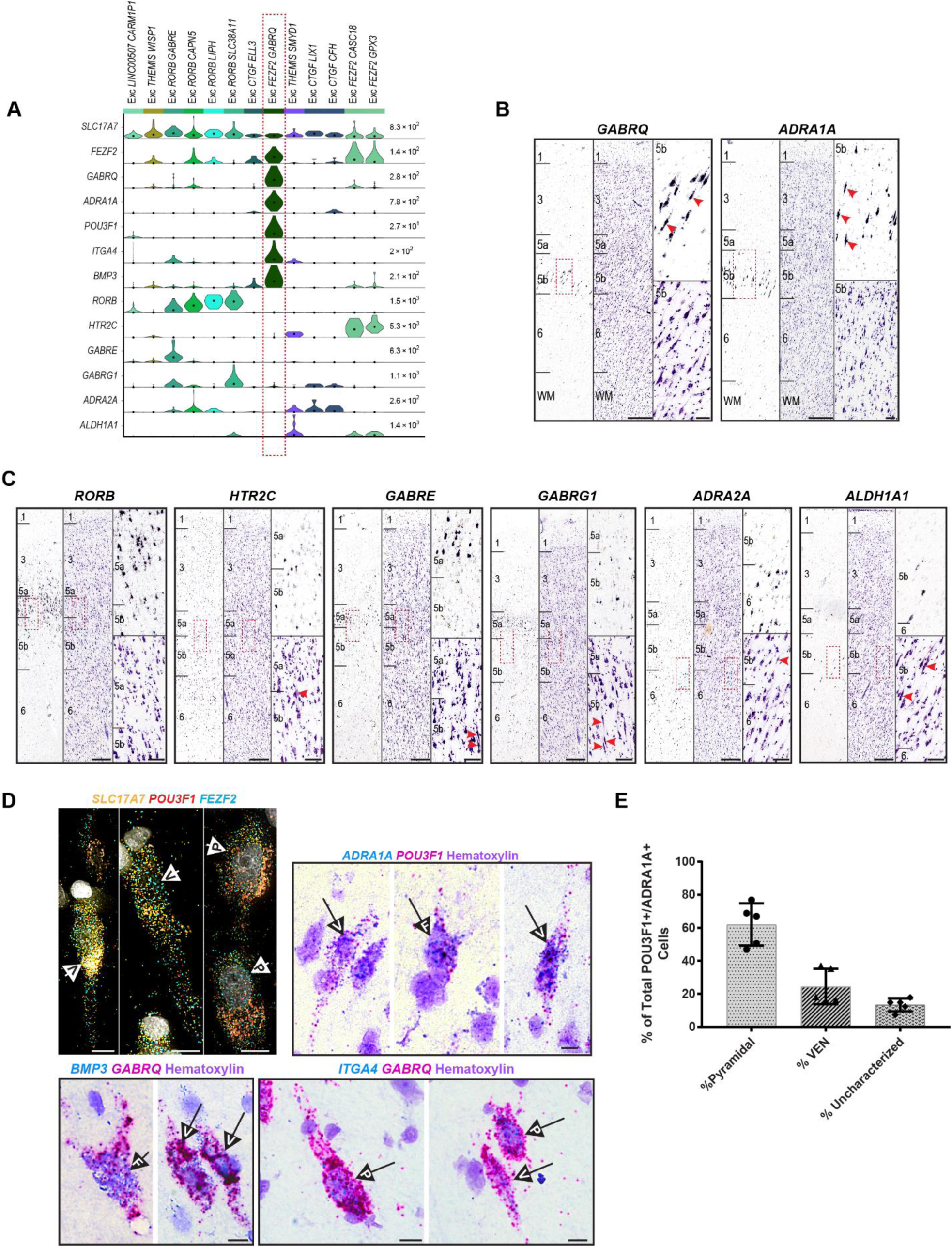
Identifying a transcriptomic cell type that corresponds to von Economo neurons (VENs) in situ. **(A)** Violin plots showing distributions of genes further examined by in situ hybridization (ISH). Each row represents a gene, black dots indicate median gene expression within clusters, and maximum gene expression values are shown on the right-hand side of each row. Gene expression values are displayed on a linear scale. The EXC GABRQ FEZF2 type expresses GABRQ and ADRA1A, previously defined markers of VENs. **(B)** Chromogenic single gene ISH from the Allen Human Brain Atlas for GABRQ and ADRA1A confirms a subset of layer 5b cells expressing these genes have spindle-shaped cell bodies typical of VENs (red arrows). The nearest Nissl-stained section is shown for each ISH image for laminar context. **(C)** ISH from the Allen Human Brain Atlas for genes expressed in other excitatory neuron types revealed by our analyses. Genes are expressed in and around layer 5 of FI but labeled cells lack spindle-shaped cell bodies typical of VENs. Red arrows in the nearest Nissl-stained section for each ISH image show cells with VEN morphology in the approximate region highlighted in the neighboring ISH image (red rectangle). Scale bars in B and C: low magnification, 150 μm, high magnification 50 μm. **(D)** Multiplex fluorescent ISH (top left) and double chromogenic ISH for marker genes of Exc FEZF2 GABRQ. Cells with pyramidal (P), VEN (V), and fork (F) morphologies are indicated by labeled arrows in each image. Scale bars, 10 μm. **(E)** Quantification of the proportion of ADRA1A+, POU3F1+ cells with pyramidal versus VEN morphologies (n=5 human donors). Cells lacking defining features of these morphological classes were called uncharacterized. Bars show the mean and error bars the standard deviation.

### Identifying a transcriptomic cell type corresponding to VENs

To characterize each cluster and determine whether one might represent VENs, we examined selective marker genes for each excitatory cluster (the top four per cluster are shown in Fig. 1d). One cluster, Exc *FEZF2 GABRQ*, specifically expressed the reported VEN and fork cell markers *GABRQ* and *ADRA1A* ^27^, suggesting that this cluster, but not the other two *FEZF2*+ clusters (Fig. 1c), likely included VENs. Exc *FEZF2 GABRQ* also had the largest number of expressed genes (**Supplementary Fig. 1b**), suggesting high RNA content and perhaps correlated with the reported large size of VENs ^1,12^. To confirm that Exc *FEZF2 GABRQ* included VENs, we looked for genes selective for one or more excitatory cell types in our dataset that also had existing ISH data in the Allen Human Brain Atlas (http://human.brain-map.org/) ^43,34^ (Fig. 2a).

As previously reported ^27^ and supporting identification of Exc *FEZF2 GABRQ* as the cluster containing VENs, ISH for *GABRQ* and *ADRA1A* showed that a subset of cells in layer 5b expressing these genes have spindle-shaped cell bodies typical of VENs (Fig. 2b). However, not all cells labeled with *GABRQ* and *ADRA1A* had spindle-shaped cell bodies, indicating that these genes do not exclusively mark VENs. In contrast, ISH for genes expressed in other excitatory neuron types did not label cells with obviously spindle-shaped cell bodies (Fig. 2c). In particular, both *HTR2C* and *ALDH1A1,* which were expressed in the other two *FEZF2*+ cell types that we found, did not label any spindle-shaped cells.

To further validate and explore the morphological cell types comprising the Exc *FEZF2 GABRQ* cluster, we performed multiplex fluorescent (mFISH) and double chromogenic (dISH) ISH for cluster-specific marker genes (Fig. 2a, d). Consistent with single gene ISH for *GABRQ* and *ADRA1A*, we find that pyramidal-shaped neurons, fork cells, and VENs are all labeled with combinations of specific marker genes for Exc *FEZF2 GABRQ* suggesting that this single transcriptomic type contains a mixture of morphological cell types (Fig. 2d). To quantify the proportions of Exc *FEZF2 GABRQ* cells with these different morphologies, we performed dISH staining for *ADRA1A+* and *POU3F1+* double-positive cells because high expression of these genes coupled with significant amplification of signal intensity in the dISH method highlights the morphology of labeled cells (Fig. 2d). Double-positive cells were classed as pyramidal, VEN, or uncharacterized (cells that lacked defining morphological features and were likely bisected by the plane of section, **Methods**) in FI tissues from 5 different human donors. Fork cells were extremely rare and were not explicitly quantified. Our results show that of all *ADRA1A*+ and *POU3F1*+ double-positive cells, ∼60% had pyramidal morphology compared with ∼25% that had VEN morphology, confirming that morphologically-defined VENs represent only a subset of the neurons that comprise the Exc *FEZF2 GABRQ* type.

### VENs are predicted to be regionally specialized extratelencephalic-projecting neurons

New methods that enable alignment of cells between data sets based on gene expression profiles can be used to align cell types across cortical regions and across species ^39,40,41^. This provides a mechanism for predicting cellular properties of human cell types based on measurements made in homologous cell types from model systems. For example, by performing retrograde labeling and scRNA-seq on the same cells (i.e. Retro-seq), the long-range projection specificity of each excitatory transcriptomic type in mouse cortex can be assessed ^32^. Previously we showed that nearly all transcriptomically-defined cell classes and subclasses identified in human MTG can be aligned with transcriptomically-defined types in mouse anterior lateral motor cortex (ALM) and primary visual cortex (VISp), even if other features are distinct between species ^33^, enabling prediction of the projection targets of homologous human types.

To shed light on the cellular properties of the Exc *FEZF2 GABRQ* type, we combined the present data from human FI with representative sets of cells from human MTG ^33^ and mouse ALM and VISp ^32^ into an integrated reference using Seurat (V3.0) ^39,40^. Only excitatory cells from each data set were included in the assembly, and cells from mouse data sets were grouped based on subclass, which combines cell types with matched predominant layer of soma location and long-range projection targets ^32^. Eight clusters were identified using Seurat, which each contain cells from all four data sets (Fig. 3a) and that matched with the groupings visualized through UMAP dimensionality reduction (Fig. 3b). More importantly, nearly all cells from mouse were mapped to the cluster in the joint assembly that matched their initially assigned subclass (Fig. 3c), with one exception. As reported in mouse ALM ^32^, we identify one cluster in agranular human FI (Exc *RORB SLC38A11*) whose best match is with intratelencephalic (IT) layer 4 clusters in human MTG and mouse VISp. Furthermore, the subclass assignments here match those reported previously using different alignment strategies for almost all MTG clusters (compare Fig. 3c with Fig 6f from ^33^). This analysis shows that Exc *FEZF2 GABRQ* co-clusters with Exc L4-5 *FEZF2 SCN4B* from human MTG and all layer 5 ET clusters from mouse VISp and ALM (Fig. 3c), suggesting that VENs are part of a cluster of neurons with deep subcortical projections.

**Figure 3.**
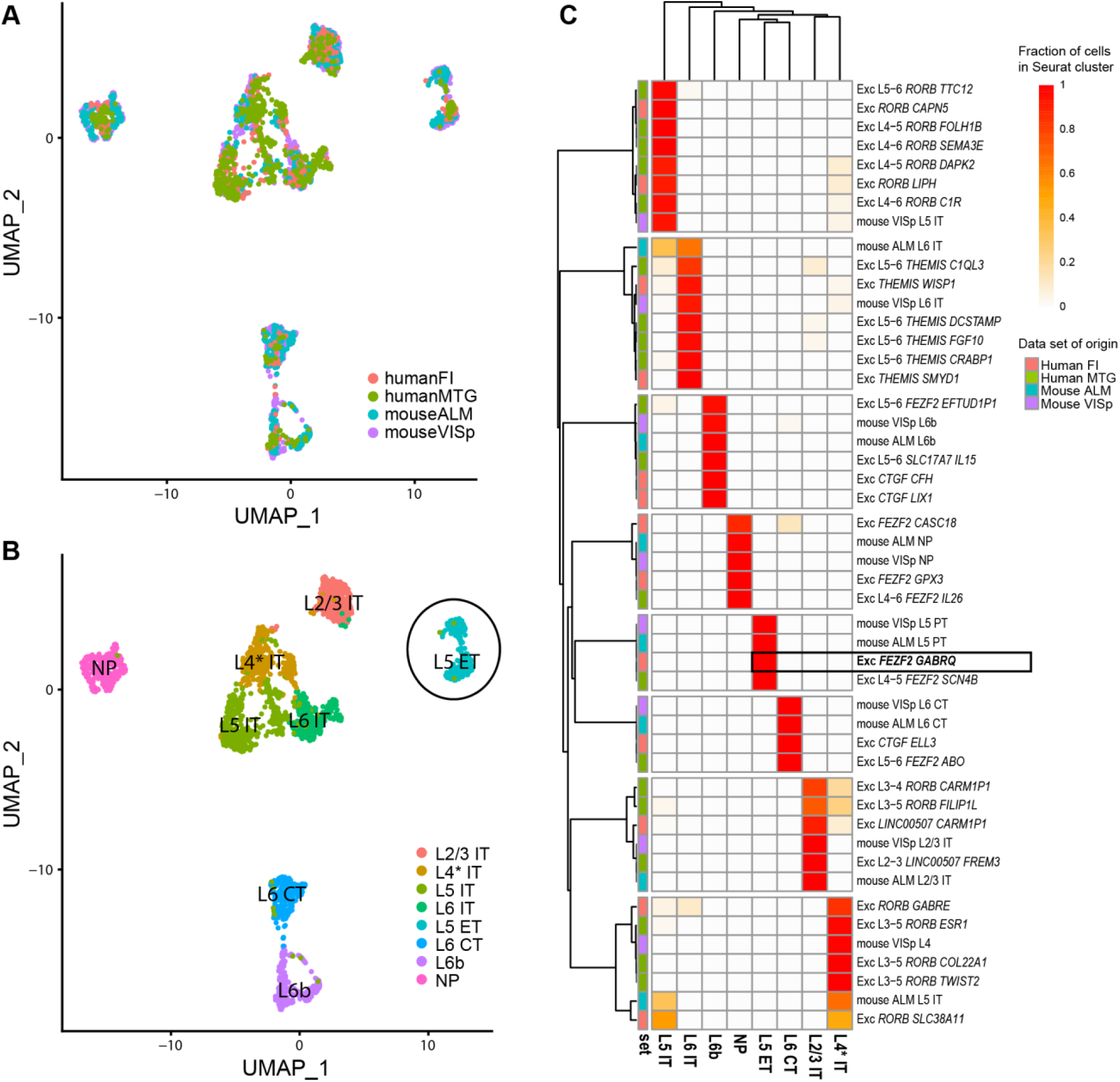
Evolutionary conservation of cell types between human and mouse predicts that VENs project sub-cortically. **(A-B)** Excitatory neurons in human FI (red), human MTG (green), mouse ALM (cyan), and mouse VISp (purple) were integrated and aligned using Seurat v3 ^37^ with default parameters, and visualized using UMAP. **(A)** Cells from each data-set co-cluster, indicating good matching of types between brain regions and species. **(B)** Eight Seurat clusters were identified using the Louvain algorithm and labeled based on expected cortical layer and projection target (as described in **C**). **(C)** Membership of cells from excitatory clusters in each data set in the Seurat clusters. Colors indicate the fraction of total cells per cluster assigned to each Seurat cluster (rows sum to 1). Data set clusters are grouped based on maximal fraction of cells in the cluster. Cortical layer of cluster inferred based on predominant cortical layer of cells from mouse and human data sets, except for the layer 4 (L4* IT) cluster which primarily includes cells from layer 5 in structures without a layer 4. Projection targets of clusters are inferred based on known projection targets of clusters in mouse ALM and VISp (IT - intratelencephalic, ET - extratelencephalic, NP - near-projecting, CT - corticothalamic). Box highlights that the Exc FEZF2 GABRQ cluster is part of a L5 ET cluster.

Interestingly, the other two *FEZF2+* clusters in FI co-cluster with near-projecting neurons in mouse, indicating that, despite the developmental role of *FEZF2* in specifying subcortical projection neurons ^30^, not all neurons that express *FEZF2* in the adult brain project subcortically. A summary of these results, including common markers between species, is shown in **Supplementary Figure 2**.

### Molecular characteristics of putative extratelencephalic neurons in human cortex

While a number of VEN marker genes have been previously described^27,26,44^, we find that although most of these genes are expressed in Exc *FEZF2 GABRQ*, very few are specific to this cluster but rather are expressed in several or many other excitatory neuron types (**Supplementary Fig. 3**). To describe a more refined set of genes selectively expressed in VENs and other putative ET neurons, we performed differential expression analysis comparing Exc *FEZF2 GABRQ* to all other excitatory clusters (**Methods**) and identified 30 genes selectively expressed in Exc *FEZF2 GABRQ* (Fig. 4a). These genes included reported markers for VENs such as *GABRQ* and *ADRA1A,* as well as many novel markers. Several genes appear to be common ET markers in mouse and human, including *FAM84B*, *POU3F1*, and *ANKRD34B* (**Supplementary Fig. 2**), although many more show divergent patterning between species than between region, as previously shown ^33^.

**Figure 4.**
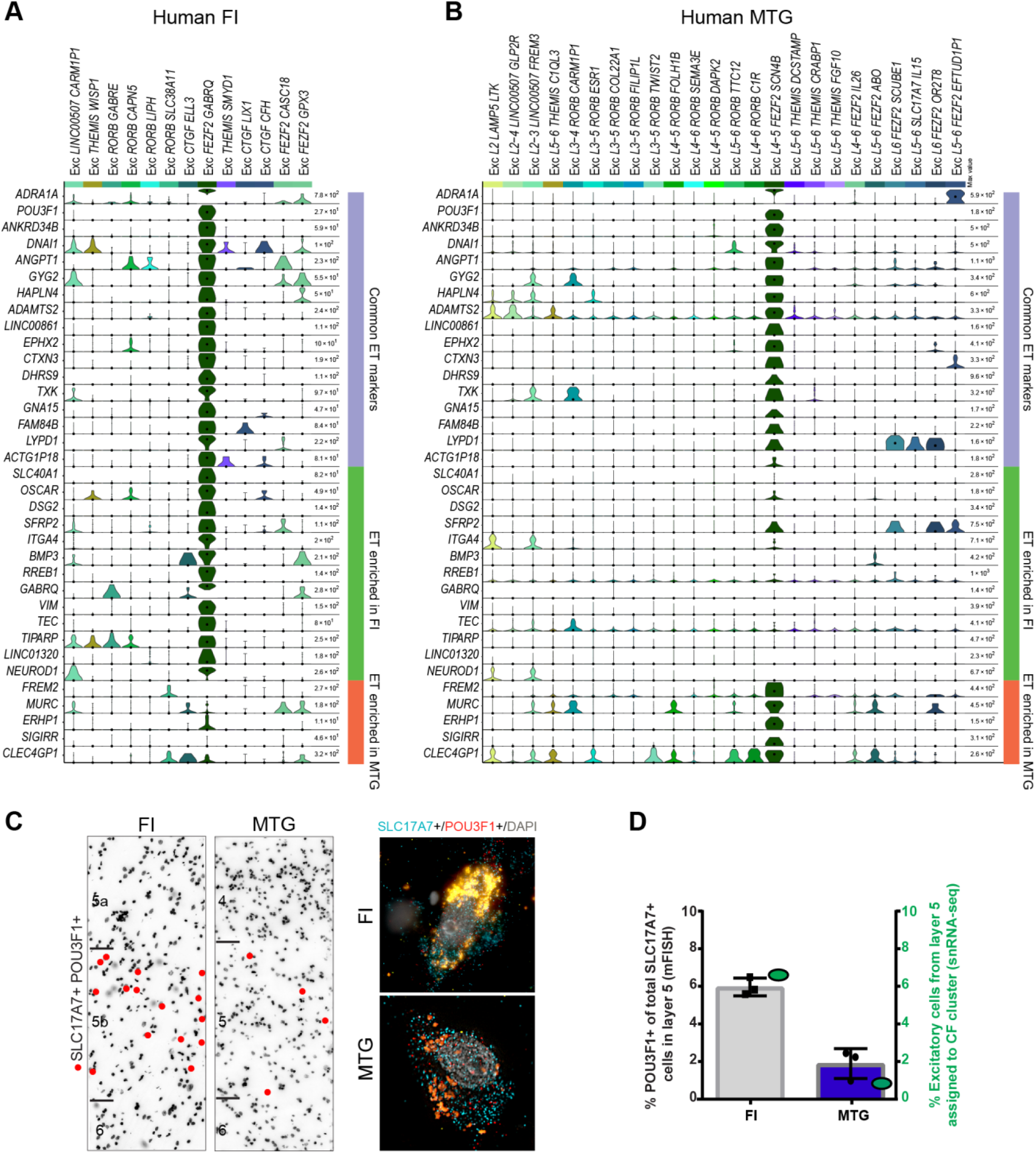
Extratelencephalic (ET) cells in human frontoinsula (FI) and middle temporal gyrus (MTG) share many common markers but differ in frequency. **(A-B)** Violin plots in human FI **(A)** and MTG **(B)** showing expression of genes enriched in Exc FEZF2 GABRQ, the corresponding cluster in human MTG, or both (see **Methods**), including many novel putative markers of VENs. Gene expression values are displayed on a log2 scale. **(C)** Representative inverted images of DAPI-stained sections of FI and MTG. Red dots depict the locations of cells labeled using multiplex fluorescent in situ hybridization (mFISH) for ET marker gene POU3F1 and SLC17A7. Scale bars in (C): DAPI images 50 μm, mFISH images 10 μm. **(D)** Black, quantification of the proportion of SLC17A7+ cells expressing the ET marker POU3F1 in FI and MTG expressed as a fraction of the total number of excitatory (SLC17A7+) cells in layer 5 of either region. Green, comparable quantification of the fraction of excitatory neurons dissected from layer 5 that are assigned to the ET clusters “Exc L4-5 FEZF2 SCN4B” (MTG) or “Exc FEZF2 GABRQ” (FI). By both mFISH and single nucleus RNA-seq (snRNA-seq), a higher fraction of putative ET cells is found in FI. Bars show the mean and error bars the standard deviation.

Interestingly, approximately half of genes enriched in Exc *FEZF2 GABRQ* were similarly enriched in the matching MTG ET cluster (e.g., *ADRA1A*) (**Methods**, Fig. 4b). However, region-specific genes were apparent for both FI (Exc *FEZF2 GABRQ*) and MTG (Exc L4-5 *FEZF2 SCN4B)*, consistent with reported variation of excitatory neurons across cortical areas ^32^, and more region-specific marker genes were apparent in FI (e.g. *GABRQ*) compared to MTG (e.g. *FREM2*). SnRNA-seq data suggested that the proportion of putative ET neurons may also vary between MTG and FI (Fig. 4c). To further examine this difference *in situ* we used mFISH to count the fraction of total excitatory cells (*SLC17A7*+) in layer 5 that also express the ET marker gene *POU3F1* (Fig. 4c, d). In agreement with snRNA-seq data, mFISH counts showed that a substantially higher fraction of putative ET cells was found in FI than in MTG. Together these results indicate that, while the primary features of putative ET neurons in human are conserved across cortical areas, ET neurons in FI appear to be more abundant and have a greater number cluster-enriched genes (Fig. 4) and more diverse cellular morphologies than those in MTG (Fig. 2, 4).

### Intrinsic membrane properties of putative VENs

ET neurons possess distinctive intrinsic membrane properties from neighboring non-ET neurons^45,46^. To test whether VENs also have distinctive electrophysiological properties, we took advantage of a very rare opportunity to perform single neuron patch clamp recordings in human insula *ex vivo* brain slices from a single human donor. In this case study, peri-tumor insula tissue was removed from the brain of a 68-year-old female patient to access a deep brain tumor located in the left insula/putamen region (Fig. 5a-c). We performed whole-cell patch clamp recordings from large spindle-shaped neurons (putative VENs) in layer 5 (n=3) and nearby (presumably non-ET) pyramidal neurons for comparison (n = 5). A biocytin cell fill was also recovered for one recorded VEN (Fig. 5d), with confirmed layer 5 localization based on soma location (1.7 mm from the pial surface of the slice) in the DAPI stain.

**Figure 5.**
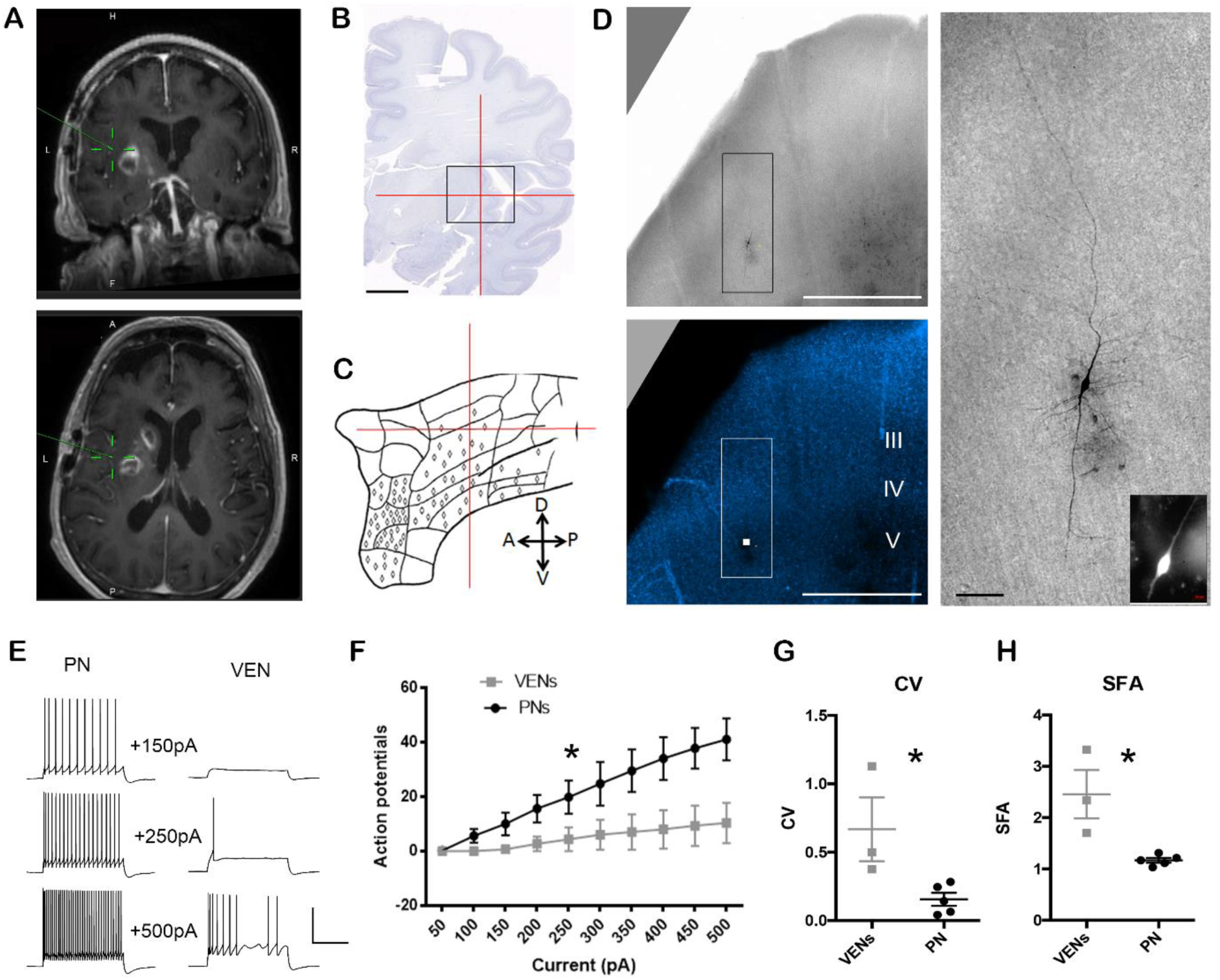
Distinctive electrophysiological properties of putative L5 VENs in ex vivo insula brain slices from a human neurosurgery patient. **(A)** MRI image data indicating the location of the excised insula tissue specimen for research. **(B)** Best matched location in the Allen 2D coronal human brain reference atlas, with crosshairs centered on the short insular gyrus. Scale bar: 1cm. **(C)** Reported distribution and relative density of VENs in the human insula (adapted from Figure 13d in Nieuwenhuys 2012 Progress in Brain Research). **(D)** Biocytin-filled putative VEN in L5 of an ex vivo insula brain slice. Low magnification brightfield and DAPI image confirms the L5 location of the neuron. The boxed region bounding the biocytin-filled neuron in expanded at right. Inset: image of Alexa dye fill following patch clamp recording in live tissue. Scale bars: 1 mm and 100 microns. **(E)** Example traces of action potential firing pattern in response to current injection steps for a representative pyramidal neuron (PN) and VEN. Scale bars: 50pA, 500msec. **(F)** Summary plot of action potential firing in response to current injection steps. *p<0.0001, 2-way ANOVA. **(G)** Summary plot of coefficient of variation (CV) for VENs versus PNs. *p<0.05, Mann-Whitney. **(H)** Summary plot of spike frequency adaptation (SFA) for VENs versus PNs. *p<0.05, Mann-Whitney.

This cell displayed the expected large spindle-shaped morphology with large caliber bipolar dendrites that extended into layer 6 (descending trunk), as well as towards the pial surface into upper layer 3 (ascending trunk). Dendritic branching was very simple, but with notable short and wispy lateral branches concentrated proximal to the soma. The axon could not be readily distinguished from these finer dendrites. The fill quality was not sufficient to make out clear dendritic spines; however, these recorded VENs appear to have a lower spine density than recorded pyramidal cells, which would be consistent with previous reports based on Golgi staining^3^.

We observed marked differences in the suprathreshold response of putative VENs versus neighboring pyramidal neurons in response to 1s current injection steps (Fig. 5e-h). Specifically, VENs produced fewer action potentials in response to a given level of current injection. This difference may be related to differences in spike timing during a train of action potentials; putative VENs displayed higher variability in spike timing and greater spike frequency accommodation than neighboring pyramidal neurons. All putative VENs displayed brief pauses and prominent subthreshold membrane oscillations during sustained firing. Although this result was not statistically significant (p>0.05), the differences in average input resistance of the putative VENs (61 ± 14 Mohms, mean ± standard error [SE]) compared with neighboring pyramids (113 ± 25 Mohm, mean ± SE) may also contribute to differences in firing of these morphologically-distinct types.

## Discussion

To determine if VENs represent a discrete transcriptionally-defined cell type, we applied snRNA-seq to classify neurons in FI layer 5 and carried out cross-species homology mapping to make predictions about VEN cellular phenotypes that are difficult to measure in human tissues. We define 13 excitatory neuron types, including one type (Exc *FEZF2 GABRQ*) that contains all VENs, but also neurons with fork and pyramidal morphologies. This approach identified many novel and selective marker genes of VENs and other excitatory neuron types that will facilitate better identification and study of these populations *in situ*. However, consistent with all published studies to date, we do not find a molecular signature that can distinguish VENs from transcriptionally similar fork or pyramidal neurons that comprise the Exc *FEZF2 GABRQ* type. One possibility is that these cells are not molecularly distinct in the adult, but rather represent a spectrum of morphologies established during development within a broader excitatory cell class. Alternatively, the current study may have lacked the power to discriminate closely related VEN and pyramidal neuron types due to the rarity of VENs relative to all excitatory neurons in layer 5 and lack of a way to specifically enrich for these cells in human. Supporting this latter idea, greater diversity of ET neurons is seen in mouse where they are more abundant and can be selectively enriched ^32,33^, including one type projecting predominantly to myelencephalon and others targeting additional subcortical areas ^32^. Further studies using higher-throughput snRNA-seq technologies will be required to definitively answer this question, but it is clear that VENs, fork cells and a subset of pyramidal cells are transcriptomically similar to one another.

Homology mapping to mouse strongly supports that VENs are deep-projecting ET neurons. Prior studies in mouse demonstrate a robust division between locally-projecting (IT) and deep-projecting (ET) neurons ^32^. Alignment of human FI data with mouse cortical scRNA-seq data shows that Exc *FEZF2 GABRQ* is highly homologous to ET neurons in mouse VISp and ALM ^32^. In addition, VENs express transcription factors required for the generation of subcortically-projecting neurons, such as *FEZF2* ^30^, but do not express transcription factors associated with corticothalamic or callosal projections ^26^. Lastly, a study in rhesus monkey proposed that VENs primarily project to distant deep brain regions, including the parabrachial nucleus of dorsolateral pons and the periaqueductal gray ^25^. Together, our findings and those of previous reports ^25,26^ strongly suggest that VENs project to deep subcortical structures. However, VEN projections might not be restricted to ET targets as the tract-tracing study above indicates that some VENs project to both ipsilateral and contralateral cortical targets, potentially including VEN populations within homologous structures of the contralateral hemisphere ^25^. These results suggest that, like rosehip neurons ^42^ and interlaminar astrocytes ^33^, VENs may represent species-specific morphological specialization of an evolutionarily shared cell type. Furthermore, we recently used homology mapping in human MTG and identified a putative ET type homologous to mouse ET types ^33^. This MTG ET type aligns to the FI ET type, and much of the ET molecular signature is shared between FI and MTG. However, many genes are expressed selectively by the FI type that are distinct from MTG. MTG has not been shown to contain VENs, and putative ET cells in MTG appear to have pyramidal neuron morphology (Fig. 4) ^33^, suggesting that VENs and fork cells might represent within-species regional specialization of a common class of large, subcortically-projecting excitatory neurons.

Selective loss of VENs and fork cells in FI and ACC has been proposed to contribute to several neuropsychiatric disorders characterized by social-emotional deficits ^20,18,19,21,47,16^. Many of these disorders show dysfunction of the salience network ^47^, which has key nodes in these same brain regions ^48^ and coordinates the brain’s responses to behaviorally-relevant stimuli ^49^, suggesting a direct link between VEN loss and dysfunction. Additionally, the salience network has functional connectivity in several subcortical areas including parts of amygdala, striatum, dorsomedial thalamus, and substantia nigra ^48^, consistent with the predicted projection targets of VENs. However, our results suggest the possibility that bvFTD and other neuropsychiatric disorders targeting FI might result from loss of ET neurons more generally, rather than exclusive loss of VENs and fork cells. The novel markers identified here for ET neurons and other excitatory types provide numerous opportunities for a refined analysis of disease-related loss of excitatory neurons. Importantly, despite the lack of VENs in rodent brains, mouse models of bvFTD are surprisingly effective at recapitulating histopathological ^50^ and behavioral ^51^ impairments reported in humans, suggesting that additional cell types are likely affected or that rodent has a homologous type to VENs that, despite different morphology, has similar circuit function.

A major challenge in understanding human brain cellular and circuit function is a paucity of tools, techniques, and tissue. However, techniques for physiological and morphological analysis using *in vitro* slice preparations and patch clamp physiology work robustly on human tissue from neurosurgical resections ^52,53,54,55,56,57,58^. Although it is exceedingly rare for tissue to be removed from regions like FI and ACC during such surgeries, the instances in which such specimens can be collected for research purposes represent rare opportunities to collect highly valuable data in the spirit of case studies in disease, where even sparse data can provide important observations and generate testable hypotheses. From the singular such specimen collected in more than three years, we demonstrate that neurons with VEN-like morphology in layer 5 of human insula can be targeted and functionally characterized. Furthermore, these putative VENs exhibit distinctive intrinsic physiological properties compared to neighboring pyramidal neurons in the same brain region. These data represent the first reported patch clamp recordings from putative VENs in the human insula, and our findings are consistent with the hypothesis that VENs represent a functionally specialized cell type, although further evidence will be necessary to establish the exact contributions of this cell type to human brain function in health and disease.

It is essential to find experimental strategies to understand the specifics of the human brain, particularly for cell types affected by disease that are not present in widely used and genetically tractable model organisms. New technological advances built on the transcriptomic approach to cell type classification promise to accelerate progress on functional analyses of human neuron types. Patch-seq allows the combination of electrophysiological, transcriptomic and morphological analysis ^59,60^, which can in principle be applied to human brain slice studies over extended time frames with recent advances culturing of human *ex vivo* brain tissue ^52^. Furthermore, novel viral tools provide cell type-specific genetic targeting in this system ^61^, and application of enhancers for ET cells, such as the *Fam84b* enhancer that labels mouse ET cells with >90% specificity ^62^, could be used to label VENs in *ex vivo* human brain tissue. Such studies can help to further refine our understanding of the defining characteristics of VENs, potentially providing information about their local connectivity and teasing out subtle gene expression differences between ET cells with spindle, fork, and pyramidal shapes.

## Methods

### Postmortem tissue donors

After obtaining permission from decedent next-of-kin, postmortem adult human brain tissue was collected by the San Diego Medical Examiner’s office and provided to the Allen Institute for Brain Science. All tissue collection was performed in accordance with the provisions of the Uniform Anatomical Gift Act described in Health and Safety Code §§ 7150, et seq., and other applicable state and federal laws and regulations. The Western Institutional Review Board reviewed tissue collection processes and determined that they did not constitute human subjects research requiring IRB review. Tissue donors were prescreened for history of neuropsychiatric disorders, neuropathology, and infectious disease (HIV, Hepatitis B, Hepatitis C), and postmortem blood samples were sent for routine serology and toxicology testing. Specimens were further screened for RNA quality and had an RNA integrity number (RIN) ≥7. Tissues used for RNA-sequencing in this study were from two control Caucasian male donors who died from cardiovascular-related issues, aged 50 (H200.1025) and 54 (H200.1030) years, as previously described ^42^.

### Tissue processing and isolation of nuclei

Whole postmortem brain specimens were processed as previously described ^42,33^. For RNA-sequencing experiments, frontoinsula (FI) was identified on slabs of interest and vibratome sectioned as described ^42,33^ (Fig. 1). Layer 5 was microdissected from vibratome sections stained with fluorescent Nissl. Mouse monoclonal anti-NeuN antibody (EMD Millipore, MAB377) was applied to nuclei preparations followed by secondary antibody staining (goat anti mouse Alexa Fluor 594, ThermoFisher), and single-nucleus sorting was carried out on a BD FACSAria Fusion instrument (BD Biosciences) using a 130 µm nozzle following a standard gating procedure as previously described (**Supplemental Fig. 1**) ^42,33^. Approximately 10% of nuclei were NeuN– negative non-neuronal nuclei. Single nuclei were sorted into 96-well PCR plates (ThermoFisher Scientific) containing 2 µL of lysis buffer (0.2% Triton-X 100, 0.2% NP-40 (Sigma Aldrich), 1 U/µL RNaseOUT (ThermoFisher Scientific), PCR-grade water (Ambion), and ERCC spike-in synthetic RNAs (Ambion). 96-well plates were snap frozen and stored at –80 °C until use. Positive controls were pools of 10 nuclei, 10 pg total RNA, and 1 pg total RNA.

### cDNA and sequencing library preparation

Single nucleus cDNA libraries were prepared using Smart-seq2 with minor modifications as previously described ^42^. Sequencing libraries were prepared using Nextera XT (Illumina) with input cDNA at 250 pg per reaction; reactions were carried out at 1/4 the volume recommended by the manufacturer with a 10 minute tagmentation step. Libraries were sequenced on a HiSeq 4000 instrument (Illumina) using 150bp paired-end reads.

### RNA-seq processing

SnRNA-seq data was processed and analyzed as previously described ^36,42^. Briefly, following demultiplexing of barcoded reads generated on the Illumina HiSeq platform, the amplification (cDNA and PCR) and sequencing primers (Illumina) and the low-quality bases were removed using Trimmomatic 0.35 software ^63^. Trimmed reads were mapped to the human reference genome, version GRCh38 (Ensembl), guided by the version 21 annotations obtained from the GENCODE repository. RSEM 1.2.31 ^64^, TOPHAT 2.1.1, and CUFFLINKS 2.2.1 ^65^ were used to quantify transcript expression at the transcriptome (exon) and whole genome (exon plus intron) levels, respectively. Software packages fastQC 0.10.1 (http://www.bioinformatics.babraham.ac.uk/projects/fastqc/), FASTX 0.0.14 (http://hannonlab.cshl.edu/fastx_toolkit/download.html), RSeQC 2.6.1 ^66^, and RNA-seq-QC 1.1.8 ^67^ were used to generate various sequence and alignment quality metrics used for classifying sample quality. A novel pipeline (SCavenger, J.M., unpublished) was created to automate execution across statistical analysis tools, integrate preformatted laboratory and clustering metrics, and calculate new statistics specific to biases identified in the single-nuclei lab and sequence preparation protocol.

### RNA-seq quality control

To remove data from low-quality samples before downstream analysis, we implemented a random forest machine-learning classification approach as previously described ^42,68^. The overall workflow for sample quality classification and filtering was to (i) establish a training set using a representative subset of samples, (ii) collect a series of 108 quality control metrics (for example, percent unique reads, percent reads surviving trimming, transcript isoform counts) spanning both the laboratory and data analysis workflows as model features, (iii) use these training data and quality control metrics to build a classification model using the random forest method, and (iv) apply the model to the entire dataset for quality classification and data filtering.

The random forest quality control model was then applied to the data and final quality Pass-Fail classifications were determined. A Pass confidence cutoff of 0.6 or greater was used to select single-nuclei data for downstream analysis. Using this random forest model applied to the entire layer 5 dataset, 78% of 1,118 single-nuclei samples passed quality control. For these Pass samples, the average number of reads after trimming was 16,715,521 ± 20,434,739, the number of ERCC transcripts detected was 41.78 ± 4.79 out of 92, and the average number of genes detected across all passing nuclei at FPKM > 1 was 5,584 ± 2,004, giving an average coverage of 2,174 reads per human gene detected. Additional summary statistics (grouped by donor or cluster) for nuclei passing QC and included in the analysis are shown in **Supplementary Figure 1**.

### Gene expression calculation

For each nucleus, expression levels were estimated based on the scaled coverage across each gene. Specifically, bam files were read into R using the readGAlignmentPairs function in the GenomicAlignments library, and genomic coverage was calculated using the coverage function in GenomicRanges ^69^. All genes in GENCODE human genome GRCh38, version 21 (Ensembl 77; 09-29-2014) were included, with gene bounds defined as the start and end locations of each unique gene specified in the gtf file (https://www.gencodegenes.org/releases/21.html). Total counts for each gene (including reads from both introns and exons) were estimated by dividing total coverage by twice the read length (150 bp, paired end). Expression levels were normalized across nuclei by calculating counts per million (CPM).

### Clustering nuclei

Nuclei and cells were grouped into transcriptomic cell types using an iterative clustering procedure as described in Boldog et al.^42^. Briefly, intronic and exonic read counts were summed, and log2-transformed expression (CPM + 1) was centered and scaled across nuclei. Differentially expressed genes were selected while accounting for gene dropouts, and principal components analysis (PCA) followed by *t*-distributed stochastic neighbor embedding (t-SNE)^70^ was used to reduce dimensionality. Nearest-neighbor distances between nuclei were calculated, and segmented linear regression (*segmented* R package) was applied to estimate the distribution breakpoint to help define the distance scale for density clustering. The statistical significance of the separation of clusters identified by density clustering was evaluated with the R package sigclust^71^, which compares the distribution of nuclei to the null hypothesis that nuclei are drawn from a single multivariate Gaussian. Iterative clustering was used to split nuclei into subclusters until the occurrence of one of four stop criteria: (i) fewer than 6 nuclei in a cluster (because it cannot be split due a minimum cluster size of 3), (ii) no significantly variable genes, (iii) no significantly variable principal components, or (iv) no significant subclusters.

To assess the robustness of clusters, the iterative clustering procedure described above was repeated 100 times for random subsamples of 80% of nuclei. A co-clustering matrix was generated that represented the proportion of clustering iterations in which each pair of nuclei was assigned to the same cluster. Average-linkage hierarchical clustering was applied to this matrix, followed by dynamic branch cutting (R package WGCNA) with cut heights ranging from 0.01 to 0.99 in steps of 0.01. A cut height resulting in 25 clusters was selected to balance cohesion (average within cluster co-clustering) and discreteness (average between cluster co-clustering) across clusters. Finally, gene markers were identified for all cluster pairs, and clusters were merged if they lacked binary markers (gene expressed in > 50% nuclei in first cluster and < 10% in second cluster) with average CPM > 1. Clusters were marked as outliers and excluded from analysis if they contained lower quality nuclei based on QC metrics or expression of mitochondrial genes.

Cluster names were defined using an automated strategy that combined molecular information (marker genes) and anatomical information (layer of dissection). Clusters were assigned to the major classes interneuron, excitatory neuron, microglia, astrocyte, oligodendrocyte precursor, or oligodendrocyte based on maximal median cluster CPM of *GAD1, SLC17A7, C3, AQP4, CSPG4,* or *OPALIN*, respectively. Clusters were then assigned a subclass marker, defined by maximal median CPM of *LAMP5, VIP, SST, PVALB, LHX6, LINC00507, RORB, THEMIS, FEZF2, CTGF, C3, FGFR3, CSPG4,* or *OPALIN.* Finally, clusters in all major classes that contained more than one cluster were assigned a cluster-specific marker gene. These marker genes had the greatest difference in the proportion of expression (CPM > 1) with a cluster compared to all other clusters regardless of mean expression level. In some cases the most specific marker gene was the subclass marker (*SST* and *VIP*).

### Scoring cluster marker genes

Many genes were expressed in the majority of nuclei in a subset of clusters. A marker score (beta) was defined for all genes to measure how binary expression was among clusters, independent of the number of clusters labeled. labeled. First, the proportion (*xi*) of nuclei in each cluster that expressed a gene above background level (CPM > 1) was calculated. Then, scores were defined as the squared differences in proportions normalized by the sum of absolute differences plus a small constant (ε) to avoid division by zero. Scores ranged from 0 to 1, and a perfectly binary marker had a score equal to 1.

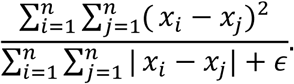

### Enrichment marker genes

Genes were defined as enriched in Exc *FEZF2 GABRQ* if they met the following criteria: 1) they were expressed in at least half the cells in Exc *FEZF2 GABRQ*, 2) they were expressed in fewer than half the cells in every other cluster, 3) they were expressed in at least 25% more cells in Exc *FEZF2 GABRQ* than in any cluster, and 4) the average expression in Exc *FEZF2 GABRQ* was at least two-fold higher than every other cluster. Thirty-two genes met these criteria.

### Cluster dendrograms

Clusters were arranged by transcriptomic similarity based on hierarchical clustering. First, the average expression level of the top 1200 scoring cluster marker genes (highest beta scores, as above) was calculated for each cluster. A correlation-based distance matrix 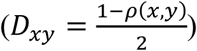 was calculated, and complete-linkage hierarchical clustering was performed using the “hclust” R function with default parameters. The resulting dendrogram branches were reordered to show inhibitory clusters followed by excitatory clusters, with larger clusters first, while retaining the tree structure. Note that this measure of cluster similarity is complementary to the co-clustering separation described above. For example, two clusters with similar gene expression patterns but a few binary marker genes may be close on the tree but highly distinct based on co-clustering.

### Gene expression visualization

Gene expression (CPM) was visualized using heat maps and violin plots, which both show genes as rows and nuclei as columns, sorted by cluster. Heat maps display each nucleus as a short vertical bar, color-coded by expression level (blue = low; red = high), and clusters were ordered as described above. The distributions of marker gene expression across nuclei in each cluster were represented as violin plots, which are density plots turned 90 degrees and reflected on the y axis. Black dots indicate the median gene expression in nuclei of a given cluster; dots above y = 0 indicate that a gene is expressed in more than half of the nuclei in that cluster.

### Colorimetric *in situ* hybridization

Information about postmortem tissue donors and methods used for colorimetric *in situ* hybridization (ISH) is available from the Allen Human Brain Atlas documentation at http://human.brain-map.org/.

### Multiplex fluorescent *in situ* hybridization (FISH)

Human tissue specimens used for RNAscope mFISH came from a cohort of neurosurgical resection and postmortem tissues that included donors used for snRNA-seq. Fresh-frozen tissues were sectioned at 14-16 μm onto Superfrost Plus glass slides (Fisher Scientific). Sections were dried for 20 minutes at −20°C and then vacuum sealed and stored at −80°C until use. The RNAscope multiplex fluorescent v1 kit was used per the manufacturer’s instructions for fresh-frozen tissue sections (ACD Bio), except that fixation was performed for 60 minutes in 4% paraformaldehyde in 1X PBS at 4°C and protease treatment was shortened to 10 minutes. Positive controls used to assess RNA quality in tissue sections were either a set from ACD Bio (*POLR2A, PPIB, UBC*, #320861) or with a combination of *SLC17A7, VIP,* and *GFAP*. Sections were imaged using either a 40X or 60X oil immersion lens on a Nikon TiE fluorescent microscope equipped with NIS-Elements Advanced Research imaging software (version 4.20). For all RNAscope mFISH experiments, positive cells were called by manually counting RNA spots for each gene. Cells were called positive for a gene if they contained ≥ 5 RNA spots for that gene. Lipofuscin autofluorescence was distinguished from RNA spot signal based on the larger size of lipofuscin granules and broad fluorescence spectrum of lipofuscin.

### Dual chromogenic *in situ* hybridization

Dual chromogenic *in situ* hybridization (dISH) was performed using the RNAscope 2.5 HD Duplex Assay Kit (ACD Bio) per the manufacturer’s protocol. Experiments were performed using fresh-frozen tissues sectioned at 16-25 μm onto Superfrost Plus glass slides (Fisher Scientific) and sections were counterstained with hematoxylin to visualize nuclei.

### Scoring of morphological types using dual chromogenic *in situ* hybridization

Staining for the EXC *FEZF2 GABRQ* markers *ADRA1A* and *POU3F1* was carried out using dISH as described above. At least 3 sections from 5 individual human donors were used for morphological assessment and scoring. First, the total number of layer 5 cells positive for both *ADRA1A* and *POU3F1* was determined for each donor. Then, the morphology of each double positive cell was assessed and scored as either pyramidal (cell body round to pyramidal in shape and wider than tall), VEN (cell body elongated, spindle-shaped and taller than wide) and uncharacterized (lacking definitive morphological features perhaps due to bisection of cells during sectioning). The proportion of cells in each morphological type was then calculated as a fraction of the total number of *ADRA1A* and *POU3F1* double positive cells. Cells were called positive for a gene if they contained ≥ 5 RNA spots for that gene.

### Electrophysiology

Electrophysiological experiments were performed as reported previously^53^. Briefly, the surgical specimen was sectioned into 300 μm thick slices using a Compresstome VF-200 (Precisionary Instruments) in a solution composed of (in mM): 92 with N-methyl-D-glucamine (NMDG), 2.5 KCl, 1.25 NaH_2_PO_4_, 30 NaHCO_3_, 20 4-(2-hydroxyethyl)-1-piperazineethanesulfonic acid (HEPES), 25 glucose, 2 thiourea, 5 Na-ascorbate, 3 Na-pyruvate, 0.5 CaCl_2_●4H_2_O and 10 MgSO_4_●7H_2_O. After warming for 10 minutes in the same solution, slices were transferred to a holding chamber containing 92 NaCl, 2.5 KCl, 1.25 NaH_2_PO_4_, 30 NaHCO_3_, 20 HEPES, 25 glucose, 2 thiourea, 5 Na-ascorbate, 3 Na-pyruvate, 2 CaCl_2_●4H_2_O and 2 MgSO_4_●7H_2_O. Slices were submerged in a recording chamber continually perfused with artificial cerebrospinal fluid (aCSF) consisting of 119 NaCl, 2.5 KCl, 1.25 NaH_2_PO4, 24 NaHCO_3_, 12.5 glucose, 2 CaCl_2_●4H_2_O and 2 MgSO4●7H_2_O and were viewed with an Olympus BX51WI microscope equipped with infrared differential contrast optics and a 40x water immersion objective.

Whole cell somatic recordings were acquired using a Multiclamp 700B amplifier and PClamp 10 data acquisition software (Molecular Devices). Electrical signals were digitized at 20-50kHz and filtered at 2-10 kHz. The pipette solution contained 130 K-gluconate, 4 KCl, 10 HEPES, 0.3 EGTA, 10 Phosphocreatine-Na_2_, 4 Mg-ATP, 0.3 Na_2_-GTP, 0.5% biocytin and .020 Alexa 594. Pipette capacitance was compensated and the bridge was balanced throughout the recording.

Data were analyzed using custom analysis scripts written in Igor Pro (Wavemetrics). All measurements were made at resting potential. FI curves were constructed by measuring the number of action potentials elicited by 1 s long current injections of increasing amplitude (Δ50 pA). Spike frequency accommodation was determined from the current injection yielding 10 ± 2 spikes and was calculated as the ratio of the last to the second interspike interval. The coefficient of variation of spike times was calculated from the same sweep.

### Quantification of putative extratelencephalic (ET) neurons

The fraction of ET neurons in FI and MTG was estimated using both mFISH and RNA-Seq. For mFISH estimates, the total numbers of *SLC17A7*+, *POU3F1*+ and *SLC17A7*+, *POU3F1*- cells in layer 5 were quantified in at least 3 sections per donor (n=3 donors for both FI and MTG). The percentage of ET cells (*SLC17A7*+, *POU3F1*+) was then calculated as a fraction of the total number of *SLC17A7*+ cells in layer 5. RNA-seq estimates were made by taking the total number of neurons mapping to the relevant ET cluster (Exc *FEZF2 GABRQ* and Exc L4-5 *FEZF2 SCN4B* in FI and MTG, respectively) and dividing by the total number of excitatory neurons collected in layer 5 dissections.

### Cross-species data integration

To assess cross-species cell type homology, excitatory cells (mouse) or nuclei (human) collected from human FI (these data), human MTG ^33^, mouse VISp, and mouse ALM ^32^ were compared. Log2-transformed CPM of intronic plus exonic reads was used as input for all four datasets. Including exonic reads increased experimental differences due to measuring whole cell versus nuclear transcripts, but this was out-weighed by improved gene detection. To the extent possible, a matched subset of cells was included as input to Seurat. In human MTG, we included all cells dissected from layers 4 or 5 that were mapped to excitatory clusters with at least 10 total cells from layer 5, including up to 50 randomly sampled cells per cluster (for a total of 616 nuclei); cells from layer 4 were included since FI does not contain a layer 4. In mouse VISp and ALM, cells were grouped by subclass (rather than cell type) and we selected 100 random cells per subclass (for a total of 700 in ALM, which does not contain layer 4, and 800 in VISp). All genes that could be matched between data sets, except a set of sex and mitochondrial genes, were considered.

These data sets were assembled into an integrated reference using Seurat V3 (https://satijalab.org/seurat/) ^39,40^ following the tutorial for Integration and Label Transfer and using default parameters for all functions, except when they differed from those used in the tutorial. More specifically, we first selected the union of the 2,000 most variable genes in each data set (using FindVariableFeatures with method=“vst”). Next, we projected this data sets into subspace based on common correlation structure using canonical correlation analysis (CCA) followed by L2 normalization, and found integration anchors (cells that are mutual nearest neighbors between data sets) in this subspace. Each anchor is weighted based on the consistency of anchors in its local neighborhood, and these anchors were then used as input to guide data integration (or batch-correction), as proposed previously ^72^. We then scaled the data, reduced the dimensionality using principal component analysis, and visualized the results with Uniform Manifold Approximation and Projection (UMAP) ^73^. We defined homologous cell types by constructing a shared nearest neighbor (SNN) graph on the integrated data sets based on the Jaccard similarity of the 10 nearest neighbors of each sample. Louvain community detection was run to identify clusters that optimized the global modularity of the partitioned graph. Data set clusters are grouped based on the maximal fraction of cells in these Seurat-assigned cluster, which were nearly perfectly aligned for most subclasses, including ET. Changes in parameters did not change the integration of cluster Exc *FEZF2 GABRQ* with mouse ET clusters.

## Data availability

Raw and aligned data have been registered with dbGaP (https://www.ncbi.nlm.nih.gov/projects/gap/cgi-bin/study.cgi?study_id=phs001791.v1.p1) and have been deposited in the NeMO archive (https://nemoarchive.org/). Specific links to these controlled-access data will be included on the dbGaP site once they become available.

## Code availability

Custom R code and count data used to generate transcriptomics related figures can be downloaded from https://github.com/AllenInstitute/L5_VEN.

## Acknowledgements

We thank Boaz Levi, Nadiya Shapovalova, and Susan Bort for assistance with flow cytometry. We acknowledge the Tissue Procurement and Tissue Processing teams at the Allen Institute for processing all neurosurgical tissues and human postmortem brain specimens. We thank Joe Davis and the San Diego Medical Examiner’s office for coordinating postmortem tissue donations. We gratefully acknowledge Nathan Hansen and our collaborators at Swedish Medical Center in Seattle for coordinating human neurosurgical tissue donations. We thank Lena Christiansen and Fan Zhang (Illumina Inc.) for assistance with sequencing. Research reported in this publication was supported by the National Institute of Neurological Disorders and Stroke of the National Institutes of Health under award number R25NS095377. The content is solely the responsibility of the authors and does not necessarily represent the official views of the National Institutes of Health. This work was supported by the Allen Institute for Brain Science, the JCVI Innovation Fund, and the Chan Zuckerberg Initiative DAF, an advised fund of the Silicon Valley Community Foundation (2018–182730). The authors thank the Allen Institute for Brain Science founder Paul G. Allen for his vision, encouragement, and support.

## Author contributions

E.S.L, R.H.S, and R.S.L conceptualized and supervised the study. R.D.H., M.N., J.T.T., B.E.K, E.R.B., M.L.B-C., F. D-F., S.I.S., K.A.S., A.M.Y., and D.N.T. contributed to experiments. J.A.M., T.E.B., B.D.A., J.M., N.J.S., P.V., and R.H.S. contributed to computational analyses. S-L.D. provided guidance on neuroanatomy. S.M.S. provided project management. A.B. and J.W.P. contributed to single-nucleus RNA-seq development. C.K. provided institutional support and project oversight. R.D.H., J.A.M., and E.S.L. wrote the manuscript with contributions from J.T.T. and B.E.K, and in consultation with all authors.

## References

1. Seeley, W. W. et al. Distinctive neurons of the anterior cingulate and frontoinsular cortex: a historical perspective. Cereb Cortex 22, 245–50 (2012).

2. Von Economo, C. A New Type of Special Cells of the Cingulate and Insular Lobes. Z Ges Neurol Psychiatr 100: 706–712, (1926).

3. Watson, K. K., Jones, T. K. & Allman, J. M. Dendritic architecture of the von Economo neurons. Neuroscience 141, 1107–12 (2006).

4. Allman, J. M. et al. The von Economo neurons in frontoinsular and anterior cingulate cortex in great apes and humans. Brain Structure and Function 214, 495–517 (2010).

5. Raghanti, M. A. et al. A Comparison of the Cortical Structure of the Bowhead Whale (Balaena mysticetus), a Basal Mysticete, with Other Cetaceans. Anat Rec (Hoboken) (2018).

6. Dijkstra, A. A., Lin, L. C., Nana, A. L., Gaus, S. E. & Seeley, W. W. Von Economo Neurons and Fork Cells: A Neurochemical Signature Linked to Monoaminergic Function. Cereb Cortex 28, 131–144 (2018).

7. Hakeem, A. Y. et al. Von Economo neurons in the elephant brain. Anat Rec (Hoboken) 292, 242–8 (2009).

8. Butti, C., Sherwood, C. C., Hakeem, A. Y., Allman, J. M. & Hof, P. R. Total number and volume of Von Economo neurons in the cerebral cortex of cetaceans. J Comp Neurol 515, 243–59 (2009).

9. Stimpson, C. D. et al. Biochemical specificity of von Economo neurons in hominoids. Am J Hum Biol 23, 22–8 (2011).

10. Allman, J. M. et al. The von Economo neurons in the frontoinsular and anterior cingulate cortex. Ann N Y Acad Sci 1225, 59–71 (2011).

11. Raghanti, M. A. et al. An analysis of von Economo neurons in the cerebral cortex of cetaceans, artiodactyls, and perissodactyls. Brain Struct Funct 220, 2303–14 (2015).

12. González-Acosta, C. A., Escobar, M. I., Casanova, M. F., Pimienta, H. J. & Buriticá, E. Von Economo Neurons in the Human Medial Frontopolar Cortex. Front Neuroanat 12, 64 (2018).

13. Raghanti, M. A. et al. An analysis of von Economo neurons in the cerebral cortex of cetaceans artiodactyls, and perissodactyls. Brain Structure and Function 220, 2303–2314 (2014).

14. Ngowyang, G. Neuere Befunde über die Gabelzellen. Zeitschrift für Zellforschung und Mikroskopische Anatomie Volume 25, Issue 2, pp 236–239, (1936).

15. Kim, E.-J. Et al. Selective Frontoinsular von Economo Neuron and Fork Cell Loss in Early Behavioral Variant Frontotemporal Dementia. Cerebral Cortex 22, 251–259 (2011).

16. Seeley, W. W. et al. Early frontotemporal dementia targets neurons unique to apes and humans. Ann Neurol 60, 660–7 (2006).

17. Nana, A. L. et al. Neurons selectively targeted in frontotemporal dementia reveal early stage TDP-43 pathobiology. Acta Neuropathol 137, 27–46 (2019).

18. Brüne, M. et al. Von Economo neuron density in the anterior cingulate cortex is reduced in early onset schizophrenia. Acta Neuropathologica 119, 771–778 (2010).

19. Brüne, M. et al. Neuroanatomical Correlates of Suicide in Psychosis: The Possible Role of von Economo Neurons. PLoS ONE 6, e20936 (2011).

20. Santos, M. et al. von Economo neurons in autism: A stereologic study of the frontoinsular cortex in children. Brain Research 1380, 206–217 (2011).

21. Kaufman, J. A. et al. Selective reduction of Von Economo neuron number in agenesis of the corpus callosum. Acta Neuropathologica 116, 479–489 (2008).

22. Gefen, T. et al. Von Economo neurons of the anterior cingulate across the lifespan and in Alzheimer’s disease. Cortex 99, 69–77 (2018).

23. Santillo, A. F. & Englund, E. Greater loss of von Economo neurons than loss of layer II and III neurons in behavioral variant frontotemporal dementia. Am J Neurodegener Dis 3, 64–71 (2014).

24. Craig, A. D. (B. How do you feel now? The anterior insula and human awareness. Nature Reviews Neuroscience 10, 59–70 (2009).

25. Evrard, H. C., Forro, T. & Logothetis, N. K. Von Economo Neurons in the Anterior Insula of the Macaque Monkey. Neuron 74, 482–489 (2012).

26. Cobos, I. & Seeley, W. W. Human von Economo Neurons Express Transcription Factors Associated with Layer V Subcerebral Projection Neurons. Cerebral Cortex 25, 213–220 (2013).

27. Dijkstra, A. A., Lin, L.-C., Nana, A. L., Gaus, S. E. & Seeley, W. W. Von Economo Neurons and Fork Cells: A Neurochemical Signature Linked to Monoaminergic Function. Cerebral Cortex 28, 131–144 (2016).

28. Yang, L. et al. Transcriptomic Landscape of von Economo Neurons in Human Anterior Cingulate Cortex Revealed by Microdissected-Cell RNA Sequencing. Cerebral Cortex 29, 838–851 (2018).

29. Allman, J. M., Watson, K. K., Tetreault, N. A. & Hakeem, A. Y. Intuition and autism: a possible role for Von Economo neurons. Trends in Cognitive Sciences 9, 367–373 (2005).

30. Rouaux, C. & Arlotta, P. Fezf2 directs the differentiation of corticofugal neurons from striatal progenitors in vivo. Nature Neuroscience 13, 1345–1347 (2010).

31. Baker, A. et al. Specialized Subpopulations of Deep-Layer Pyramidal Neurons in the Neocortex: Bridging Cellular Properties to Functional Consequences. The Journal of Neuroscience 38, 5441–5455 (2018).

32. Tasic, B. et al. Shared and distinct transcriptomic cell types across neocortical areas. Nature 563, 72–78 (2018).

33. Hodge, R. D. et al. Conserved cell types with divergent features between human and mouse cortex. (2018). doi:10.1101/384826

34. Zeng, H. et al. Large-Scale Cellular-Resolution Gene Profiling in Human Neocortex Reveals Species-Specific Molecular Signatures. Cell 149, 483–496 (2012).

35. Grindberg, R. V. et al. RNA-sequencing from single nuclei. Proceedings of the National Academy of Sciences 110, 19802–19807 (2013).

36. Krishnaswami, S. R. et al. Using single nuclei for RNA-seq to capture the transcriptome of postmortem neurons. Nat Protoc 11, 499–524 (2016).

37. Zeisel, A. et al. Cell types in the mouse cortex and hippocampus revealed by single-cell RNA-seq. Science 347, 1138–1142 (2015).

38. Tasic, B. et al. Adult mouse cortical cell taxonomy revealed by single cell transcriptomics. Nature Neuroscience 19, 335–346 (2016).

39. Stuart, T. et al. Comprehensive integration of single cell data. (2018). doi:10.1101/460147

40. Butler, A., Hoffman, P., Smibert, P., Papalexi, E. & Satija, R. Integrating single-cell transcriptomic data across different conditions technologies, and species. Nature Biotechnology 36, 411–420 (2018).

41. Johansen, N. & Quon, G. scAlign: a tool for alignment integration and rare cell identification from scRNA-seq data. (2018). doi:10.1101/504944

42. Boldog, E. et al. Transcriptomic and morphophysiological evidence for a specialized human cortical GABAergic cell type. Nature Neuroscience 21, 1185–1195 (2018).

43. Hawrylycz, M. J. et al. An anatomically comprehensive atlas of the adult human brain transcriptome. Nature 489, 391–399 (2012).

44. Yang, L. et al. Transcriptomic Landscape of von Economo Neurons in Human Anterior Cingulate Cortex Revealed by Microdissected-Cell RNA Sequencing. Cereb Cortex 29, 838–851 (2019).

45. Baker, A. et al. Specialized Subpopulations of Deep-Layer Pyramidal Neurons in the Neocortex: Bridging Cellular Properties to Functional Consequences. J Neurosci 38, 5441–5455 (2018).

46. Gouwens, N. W. et al. Classification of electrophysiological and morphological types in mouse visual cortex. (2018). doi:10.1101/368456

47. Zhou, J. et al. Divergent network connectivity changes in behavioural variant frontotemporal dementia and Alzheimer’s disease. Brain 133, 1352–1367 (2010).

48. Seeley, W. W. et al. Dissociable Intrinsic Connectivity Networks for Salience Processing and Executive Control. Journal of Neuroscience 27, 2349–2356 (2007).

49. Menon, V. & Uddin, L. Q. Saliency switching, attention and control: a network model of insula function. Brain Structure and Function 214, 655–667 (2010).

50. Roberson, E. D. Mouse models of frontotemporal dementia. Annals of Neurology 72, 837–849 (2012).

51. Vernay, A., Sellal, F. & René, F. Evaluating Behavior in Mouse Models of the Behavioral Variant of Frontotemporal Dementia: Which Test for Which Symptom? Neurodegenerative Diseases 16, 127–139 (2015).

52. Ting, J. T. et al. A robust ex vivo experimental platform for molecular-genetic dissection of adult human neocortical cell types and circuits. Scientific Reports 8, (2018).

53. Kalmbach, B. E. et al. h-Channels Contribute to Divergent Intrinsic Membrane Properties of Supragranular Pyramidal Neurons in Human versus Mouse Cerebral Cortex. Neuron 100, 1194–1208.e5 (2018).

54. Boldog, E. et al. Transcriptomic and morphophysiological evidence for a specialized human cortical GABAergic cell type. Nat Neurosci 21, 1185–1195 (2018).

55. Beaulieu-Laroche, L. et al. Enhanced Dendritic Compartmentalization in Human Cortical Neurons. Cell 175, 643–651.e14 (2018).

56. Mohan, H. et al. Dendritic and Axonal Architecture of Individual Pyramidal Neurons across Layers of Adult Human Neocortex. Cerebral Cortex 25, 4839–4853 (2015).

57. Szabadics, J. et al. Excitatory effect of GABAergic axo-axonic cells in cortical microcircuits. Science 311, 233–5 (2006).

58. Verhoog, M. B. et al. Mechanisms underlying the rules for associative plasticity at adult human neocortical synapses. J Neurosci 33, 17197–208 (2013).

59. Cadwell, C. R. et al. Electrophysiological transcriptomic and morphologic profiling of single neurons using Patch-seq. Nature Biotechnology 34, 199–203 (2015).

60. Fuzik, J. et al. Integration of electrophysiological recordings with single-cell RNA-seq data identifies neuronal subtypes. Nat Biotechnol 34, 175–183 (2016).

61. Mich, J. K. et al. Epigenetic landscape and AAV targeting of human neocortical cell classes. (2019). doi:10.1101/555318

62. Graybuck, L. T. et al. Prospective brain-wide labeling of neuronal subclasses with enhancer-driven AAVs. (2019). doi:10.1101/525014

63. Bolger, A. M., Lohse, M. & Usadel, B. Trimmomatic: a flexible trimmer for Illumina sequence data. Bioinformatics 30, 2114–2120 (2014).

64. Li, B. & Dewey, C. RSEM. in Bioinformatics 41–74 (Apple Academic Press, 2014). doi:10.1201/b16589-5

65. Trapnell, C. et al. Differential gene and transcript expression analysis of RNA-seq experiments with TopHat and Cufflinks. Nature Protocols 7, 562–578 (2012).

66. Wang, L., Wang, S. & Li, W. RSeQC: quality control of RNA-seq experiments. Bioinformatics 28, 2184–2185 (2012).

67. DeLuca, D. S. et al. RNA-SeQC: RNA-seq metrics for quality control and process optimization. Bioinformatics 28, 1530–1532 (2012).

68. Aevermann, B. et al. Production of a preliminary quality control pipeline for single nuclei rna-seq and its application in the analysis of cell type diversity of post-mortem human brain neocortex. Pac Symp Biocomput 22, 564–575 (2017).

69. Lawrence, M. et al. Software for computing and annotating genomic ranges. PLoS Comput Biol 9, e1003118 (2013).

70. Van Der Maaten, L. J. P. & Hinton, G. E. Visualizing high-dimensional data using t-sne. Journal of Machine Learning Research 9, 2579–2605 (2008).

71. Liu, Y., Hayes, D. N., Nobel, A. & Marron, J. S. Statistical Significance of Clustering for High-Dimension, Low–Sample Size Data. Journal of the American Statistical Association 103, 1281–1293 (2008).

72. Haghverdi, L., Lun, A. T. L., Morgan, M. D. & Marioni, J. C. Batch effects in single-cell RNA-sequencing data are corrected by matching mutual nearest neighbors. Nature Biotechnology 36, 421–427 (2018).

73. Leland McInnes, J. M., John Healy. UMAP: Uniform Manifold Approximation and Projection for Dimension Reduction. *arXiv* 1802.03426, (2018).

