## Supplementary Information for "Transcriptomic evidence that von Economo neurons are regionally specialized extratelencephalic-projecting excitatory neurons"

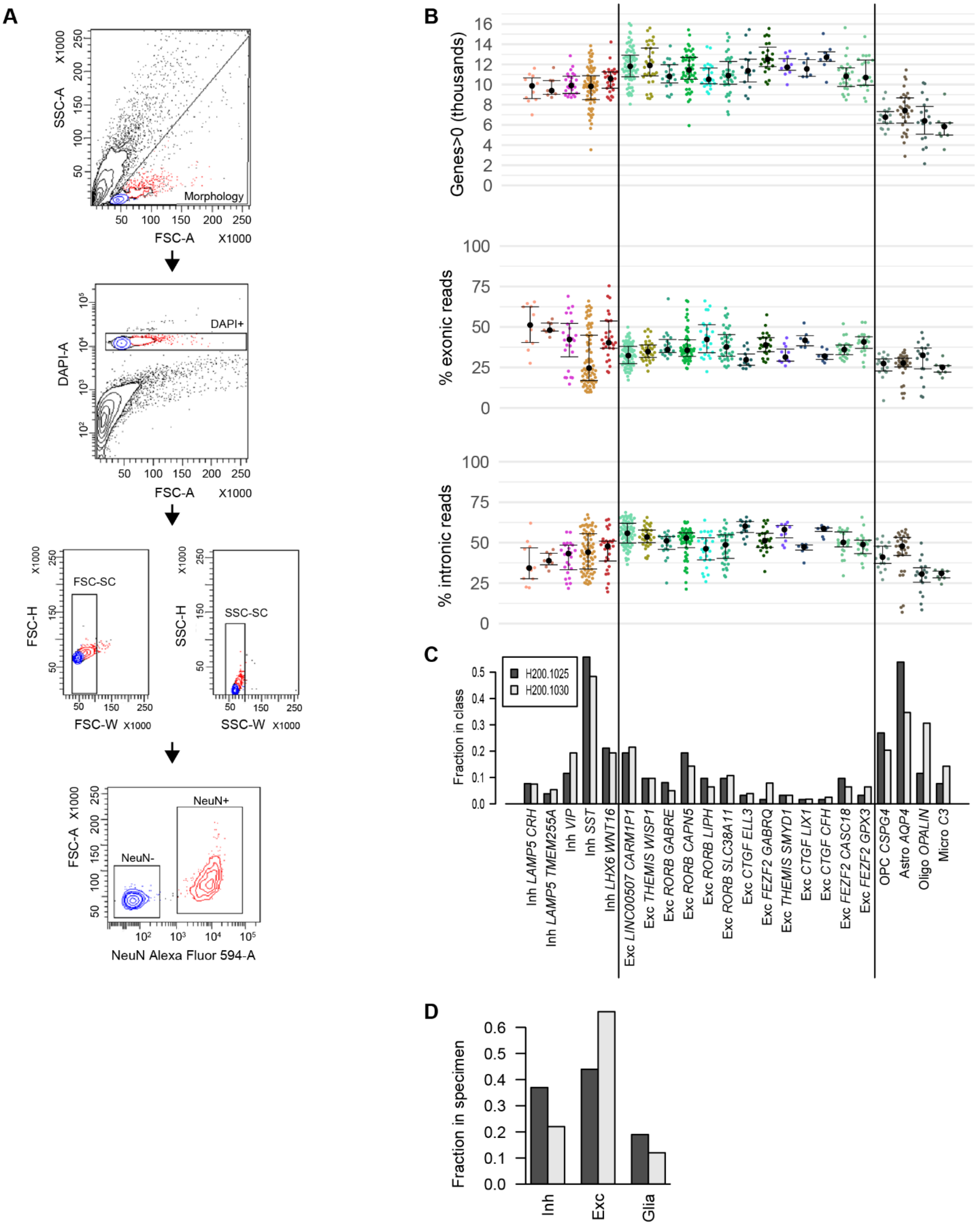

A
SSC-A X1000
FSC-A X1000
Morphology
DAPI-A
DAPI+
FSC-H X1000
FSC-W X1000
FSC-SC
SSC-H X1000
SSC-W X1000
SSC-SC
FSC-A X1000
NeuN Alexa Fluor 594-A
NeuN-
NeuN+
B
Genes>0 (thousands)
% exonic reads
% intronic reads
C
Fraction in class
H200.1025
H200.1030
Inh LAMP5 CRH
Inh LAMP5 TMEM255A
Inh VIP
Inh SST
Inh LHX6 WNT16
Exc LINC00507 CARM1P1
Exc THEMIS WISP1
Exc RORB GABRE
Exc RORB CAPN5
Exc RORB LIPH
Exc RORB SLC38A11
Exc CTGF ELL3
Exc FEZF2 GABRQ
Exc THEMIS SMYD1
Exc CTGF LIX1
Exc CTGF CFH
Exc FEZF2 CASC18
Exc FEZF2 GPX3
OPC CSPG4
Astro AQP4
Oligo OPALIN
Micro C3
D
Fraction in specimen
Inh
Exc
Glia

**Supplementary Figure 1. (A)** Standard fluorescence activated-cell sorting (FACS) gating scheme for collection of NeuN-positive and NeuN-negative nuclei. **(B)** Scatter plots plus median and interquartile interval of three QC metrics grouped and colored by cluster. Median gene detection was highest among excitatory neuron types (with Exc *FEZF2 GABRQ* the highest), lower among inhibitory neuron types, and significantly lower among non-neuronal types. All clusters had a large fraction of both exonic and intronic reads, with high cell type dependencies on these measures. **(C-D)** Bar plots showing that the fraction of nuclei from each cell type is relatively consistent between the two donors, when scaled by the total number of inhibitory, excitatory, and non-neuronal nuclei collected from each donor, but that a disproportionately smaller fraction of excitatory nuclei collected from donor H200.1025 passed QC compared both with donor H200.1030 and with expectations. Vertical lines separate major cell classes.

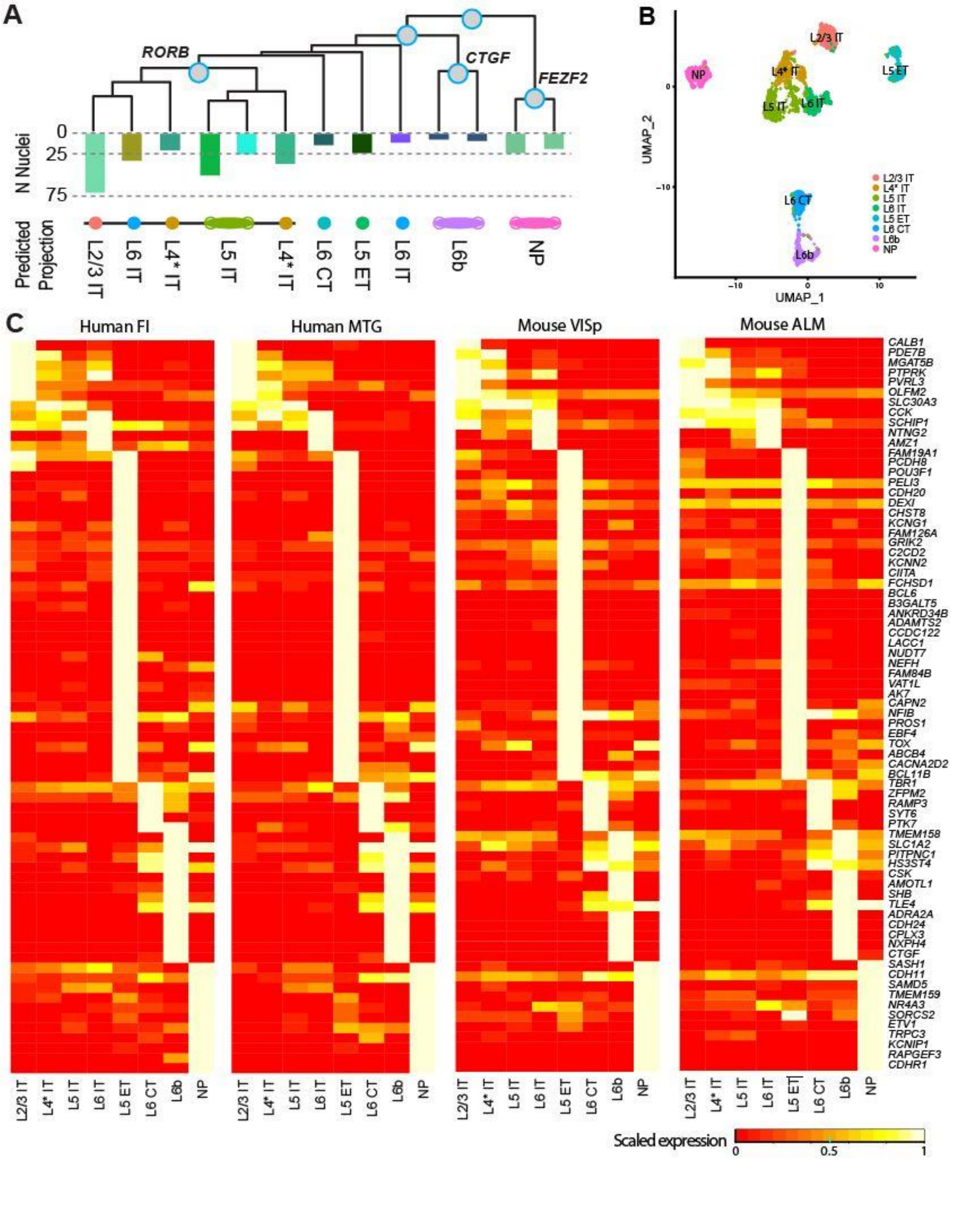

A
RORB
CTGF
FEZF2
N Nuclei
0
25
75
Predicted Projection
L2/3 IT
L6 IT
L4* IT
L5 IT
L4* IT
L6 CT
L5 ET
L6 IT
L6b
NP
B
UMAP_1
UMAP_2
L2/3 IT
L4* IT
L5 IT
L6 IT
L5 ET
L6 CT
L6b
NP
C
Human FI
Human MTG
Mouse VISp
Mouse ALM
CALB1
PDE7B
MGAT5B
PTPRK
PVRL3
OLFM2
SLC30A3
CCK
SCHIP1
NTNG2
AMZ1
FAM19A1
PCDH8
POU3F1
PELI3
CDH20
DEXI
CHST8
KCNG1
FAM126A
GRIK2
C2CD2
KCNN2
CIITA
FCHSD1
BCL6
B3GALT5
ANKRD34B
ADAMTS2
CCDC122
LACC1
NUDT7
NEFH
FAM84B
VAT1L
AK7
CAPN2
NFIB
PROS1
EBF4
TOX
ABCB4
CACNA2D2
BCL11B
TBR1
ZFPM2
RAMP3
SYT6
PTK7
TMEM158
SLC1A2
PITPNC1
HS3ST4
CSK
AMOTL1
SHB
TLE4
ADRA2A
CDH24
CPLX3
NXPH4
CTGF
SASH1
CDH11
SAMD5
TMEM159
NR4A3
SORCS2
ETV1
TRPC3
KCNIP1
RAPGEF3
CDHR1
Scaled expression
0
0.5
1

**Supplementary Figure 2. Annotation of human excitatory types using mouse**. **(A)** Eight Seurat clusters labeled based on expected cortical layer and projection target, as shown in **Figure 3B**. **(B)** Hierarchical representation of 13 excitatory cell types, as shown in **Figure 1B**, additionally labeled with their predicted projections from mouse (see **Figure 3**). **(C)** Heatmap showing 85 common marker genes for the eight matched clusters across all four data sets. These genes had high Pearson correlation of average Seurat cluster expression for all pairs of data sets ($R>0.75$) and high specificity, defined as the cumulative sum of the sorted average Seurat cluster expression, in each data set ($S<3$).

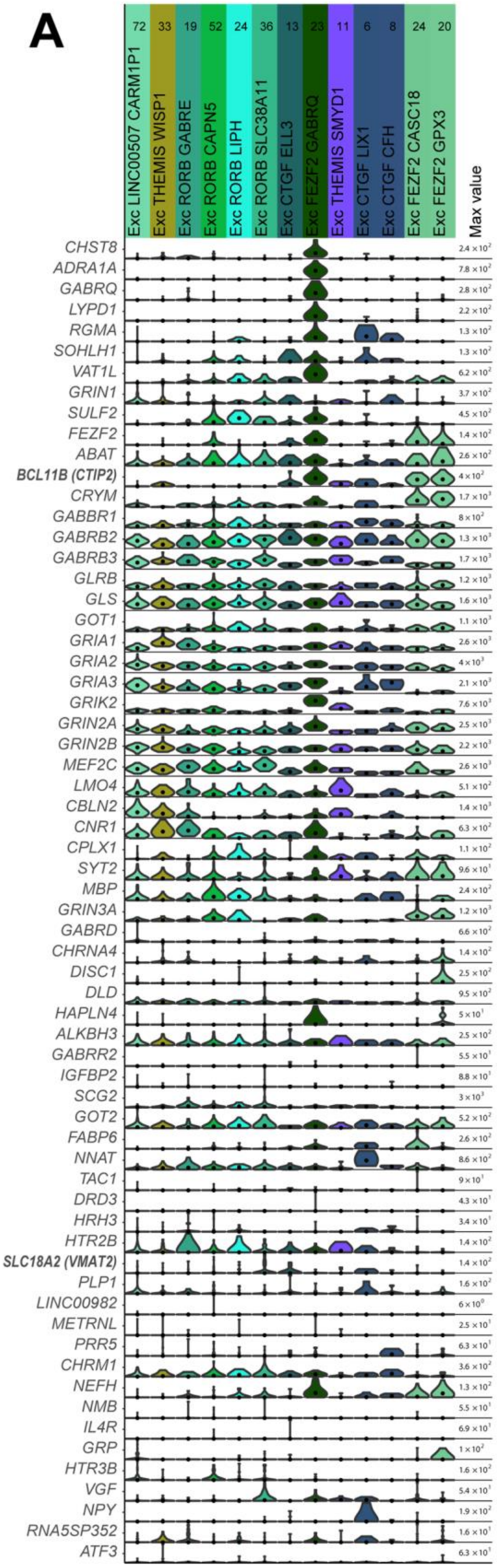


**B**

| Gene | Allman 2005 (ICC) | Allman 2010 (IHC) | Stimpson 2011 (ICC) | Cobos 2013 (ISH/IHC/dFISH) | Dijkstra 2016 (ISH from ABA, IHC/ISH*) | Yang 2018 (LMD + RNA-seq, IHC/ISH/IF*) | |
|---|---|---|---|---|---|---|---|
| *CHST8* | | | | | | X* | * |
| *ADRA1A* | | | | | X* | | * |
| *GABRQ* | | | | | X* | | * |
| *LYPD1* | | | | | | X* | * |
| *RGMA* | | | | | | X | |
| *SOHLH1* | | | | | | X | |
| *VAT1L* | | | | | | X* | * |
| *GRIN1* | | | | | e | | * |
| *SULF2* | | | | | | X* | |
| *FEZF2* | | | | X | | X | |
| *ABAT* | | | | | e | | * |
| *BCL11B* | | | | X | | | |
| *CRYM* | | | | | e | | * |
| *GABBR1* | | | | | e | | * |
| *GABRB2* | | | | | e | | * |
| *GABRB3* | | | | | e | | * |
| *GLRB* | | | | | e | | * |
| *GLS* | | | | | e | | * |
| *GOT1* | | | | | e | | * |
| *GRIA1* | | | | | e | | * |
| *GRIA2* | | | | | e | | * |
| *GRIA3* | | | | | e | | * |
| *GRIK2* | | | | | e | | * |
| *GRIN2A* | | | | | e | | * |
| *GRIN2B* | | | | | e | | * |
| *MEF2C* | | | | | | X | |
| *LMO4* | | | | e | | | * |
| *CBLN2* | | | | | e | | * |
| *CNR1* | | | | | e | | * |
| *CPLX1* | | | | | | X | |
| *SYT2* | | | | | e | | * |
| *MBP* | | | | | | X | |
| *GRIN3A* | | | | | e | | |
| *GABRD* | | | | | e | | |
| *CHRNA4* | | | | | e | | |
| *DISC1* | | X | | | | | |
| *DLD* | | | | | e | | |
| *HAPLN4* | | | | | | X | |
| *ALKBH3* | | | | | | X | |
| *GABRR2* | | | | | e | | |
| *IGFBP2* | | | | | | X | |
| *SCG2* | | | | | e | | |
| *GOT2* | | | | | e | | * |
| *FABP6* | | | | | | X | |
| *NNAT* | | | | | e | | |
| *TAC1* | | | | | e | | |
| *DRD3* | X | | | | | | |
| *HRH3* | | | | | e | | |
| *HTR2B* | X | | | | | | |
| *SLC18A2* | | | | | X* | | |
| *PLP1* | | | | | | X | |
| *LINC00982* | | | | | | X | |
| *METRNL* | | | | | | X | |
| *PRR5* | | | | | | X | |
| *CHRM1* | | | | | e | | |
| *NEFH* | | | | | e | | * |
| *NMB* | | X | X | | | | |
| *IL4R* | | | X | | | | |
| *GRP* | | X | | | | | |
| *HTR3B* | | | | | e | | |
| *VGF* | | | | | e | | |
| *NPY* | | | | | | X | |
| *RNA5SP352* | | | | | | X | |
| *ATF3* | | | X | | | | |

**Supplementary Figure 3. Expression of previously-reported VEN markers in human FI**. **(A)** Violin plots showing expression of previously reported VEN marker genes per excitatory cluster.  Around half of these genes are highly expressed in Exc *FEZF2 GABRQ*, but with highly varying levels of specificity.  Genes are roughly ordered from most to least selective for Exc *FEZF2 GABRQ* in this study.  Each row represents a gene, black dots show median gene expression within clusters, and the maximum expression value for each gene is shown on the right-hand side of each row. Gene expression values are displayed on a linear scale.  **(B)** Specific evidence from previous publications about the specificity of the genes in **A**. Each column corresponds to the listed publication, along with the methods used for identifying VEN markers (ICC, immunocytochemistry; IHC, immunohistochemistry; ISH, in situ hybridization; dFISH, double fluorescent ISH; ABA, Allen Brain Atlas; LMD, laser microdissection; IF, immunofluorescence). Marked boxes indicate that a given gene was found to be expressed in VENs in a given study (e = expressed in VENs, but not selective; X = selective in VENs as compared to surrounding pyramidal neurons; X* = validated as selective using at least one additional method, as indicated).  Note that genes validated with multiple methods or in multiple studies tend to also be the most selective for Exc *FEZF2 GABRQ* in this study. Green asterisks indicate genes showing qualitative agreement between current and previous results (e.g. specific to the VEN cluster or expressed in the VEN cluster as previously reported).
